# First annotated draft genomes of non-marine ostracods (Ostracoda, Crustacea) with different reproductive modes

**DOI:** 10.1101/2020.12.02.409169

**Authors:** Patrick Tran Van, Yoann Anselmetti, Jens Bast, Zoé Dumas, Nicolas Galtier, Kamil S. Jaron, Koen Martens, Darren J. Parker, Marc Robinson-Rechavi, Tanja Schwander, Paul Simion, Isa Schön

**Affiliations:** Department of Ecology and Evolution, University of Lausanne, 1015 Lausanne, Switzerland; Swiss Institute of Bioinformatics, Lausanne, Switzerland; ISEM – Institut des Sciences de l’Evolution, Montpellier, France; Current address: CoBIUS lab, Department of Computer Science, University of Sherbrooke, 2500 Boulevard de l’Université, Sherbrooke, QC J1K 2R1, Canada; Current address: Institute for Zoology, University of Cologne, Köln, Germany; Royal Belgian Institute of Natural Sciences, OD Nature, Freshwater Biology, Brussels, Belgium; University of Ghent, Dept Biology, Ghent, Belgium; Université de Namur, LEGE, URBE, Namur, 5000, Belgium; University of Hasselt, Research Group Zoology, Diepenbeek, Belgium

**Keywords:** ancient asexual, sexual, *Darwinula stevensoni*, *Cyprideis torosa*

## Abstract

Ostracods are one of the oldest crustacean groups with an excellent fossil record and high importance for phylogenetic analyses but genome resources for this class are still lacking. We have successfully assembled and annotated the first reference genomes for three species of non-marine ostracods; two with obligate sexual reproduction (*Cyprideis torosa* and *Notodromas monacha*) and the putative ancient asexual *Darwinula stevensoni*. This kind of genomic research has so far been impeded by the small size of most ostracods and the absence of genetic resources such as linkage maps or BAC libraries that were available for other crustaceans. For genome assembly, we used an Illumina-based sequencing technology, resulting in assemblies of similar sizes for the three species (335-382Mb) and with scaffold numbers and their N50 (19-56 kb) in the same orders of magnitude. Gene annotations were guided by transcriptome data from each species. The three assemblies are relatively complete with BUSCO scores of 92-96%, and thus exceed the quality of several other published crustacean genomes obtained with similar techniques. The number of predicted genes (13,771-17,776) is in the same range as Branchiopoda genomes but lower than in most malacostracan genomes. These three reference genomes from non-marine ostracods provide the urgently needed basis to further develop ostracods as models for evolutionary and ecological research.

## BACKGROUND

### Relevance of ostracods

Ostracoda are small, bivalved crustaceans, widely occurring in almost all aquatic habitats as part of the meiobenthos and periphyton. There are 2,330 formally described species of extant non-marine ostracods (Meisch *et al*. 2019) and at least another 7,000 described species of extant marine ostracod species (see Schön and Martens 2016 for an estimate by S. Brandao). Their calcified valves are preserved as microfossils, making them the extant arthropod group with the most extensive fossil record. The group has an estimated (Cambrian) age of c 500 myr (millions of years) according to a molecular clock (Oakley *et al*. 2013), and c. 450 myr (Ordovician; Maddocks 1982) to 509 myr (Wolfe *et al*. 2016) according to the fossil record. This makes them one of the oldest extant pancrustacean groups (Figure 1). Because of their excellent fossil data, evolutionary events can be dated with real time estimates making ostracods ideal models for evolutionary research (Butlin and Menozzi 2000; Oakley and Cunningham 2002; Oakley *et al*. 2013; Schön and Martens 2016;).

**Figure 1:**
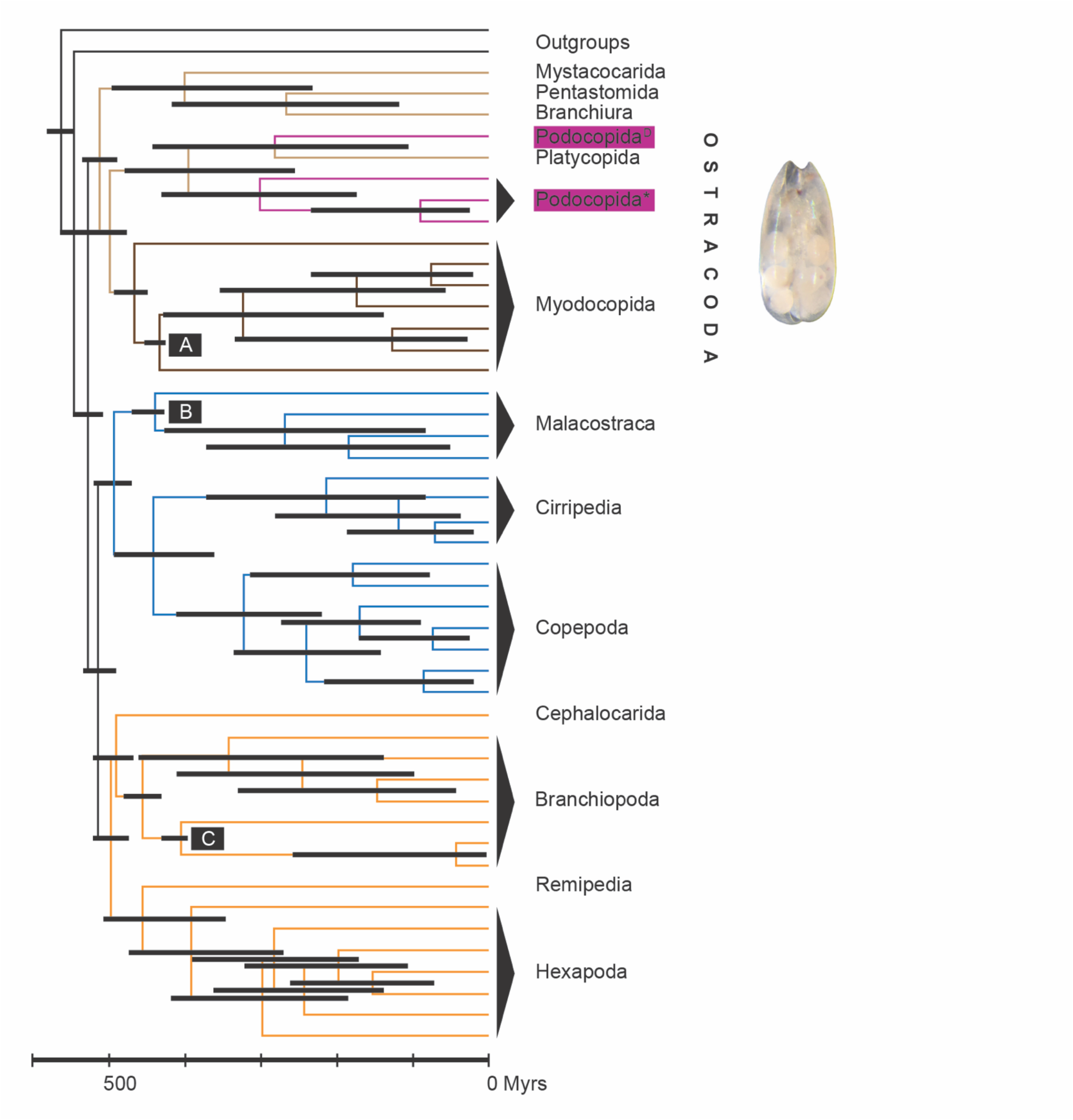
The phylogenetic position of the Ostracoda among the pancrustaceans and their age estimated from fossil and molecular data. Modified from Oakley et al. (2013). Different pancrustaceans are indicated by branches in different colours. The Ostracoda include the Podocopida, Platycopida and Myodocopida. Here, three representatives of the Platycopida (indicated in purple) have been sequenced. The phylogenetic clade to which *Darwinula stevensoni* belongs, is indicated by ^D^, the clade to which Cyprideis torosa and Notodromas monacha belong, is indicated by *. Black horizontal bars represent the range of age estimates in myr from Bayesian analyses by Oakley et al. (2013). The letters A-C in the black boxes indicated fossils that were used for calibrations of age estimates.

Contrary to the extensive focus on this group for palaeontological research, there is a total lack of published ostracod genomes, and even isolated genomic data from ostracods in open access databases are still rare. Thus, the only resources available beyond individual gene sequences are four mitogenomes (the marine ostracods *Vargula hilgendorfii* (Ogoh and Ohmiya 2004; GenBank accession number NC_005306) and *Cypridina dentata* (Wang *et al*. 2019; NC_042792); and two unpublished mitogenomes from *V. tsujii* (NC_039175) and *Cyprideis torosa* (PRJNA302529)). Also, raw Illumina DNA sequencing reads of the podocopid ostracod *Eucypris virens* have been generated as part of a study testing DNA extraction methods for high throughput sequencing in zooplankton (SRX8021019; Beninde *et al*. 2020) but these have neither been assembled nor annotated. In studies on crustacean phylogenies and gene expression (see Table S1 for details), raw RNA-sequencing reads have been generated for a total of 12 species coming from the three major ostracod lineages (Mydocopida, Halocyprida and Podocopida), but the number of assembled and annotated ostracod genes in these studies remains very limited, ranging between 4 and 822 genes.

### Choice of model species

Extant non-marine ostracods show a high prevalence of asexual reproduction (Chaplin *et al*. 1994; Butlin *et al*. 1998; Martens *et al*. 1998), which has evolved several times independently in different ostracod lineages and is most frequent in the Cyprididae and the Darwinulidae. Ostracods are thus an ideal group to further study the paradox of sex, which remains one of the most puzzling questions in evolutionary biology (Bell 1982; Otto and Lenormand 2002; Schön *et al*. 2009a; Neiman *et al*. 2018). The most important sets of hypotheses explaining why sex is advantageous despite its direct costs are based on the fact that physical linkage among loci generates different forms of selective interference (recently reviewed in Otto 2020). Genome-wide data are very valuable to test if asexuals indeed are affected by these predictions (e.g., Glémin *et al*. 2019; Jaron *et al*. 2020) and to develop insights into mechanisms such as gene conversion (Omilian *et al*. 2006), DNA repair (Schön and Martens, 1998; Hecox-Lea and Mark Welch 2018) or horizontal gene transfer (Gladyshev *et al*. 2008; Danchin *et al*. 2010; Boschetti *et al*. 2012; Flot *et al*. 2013; Paganini *et al*. 2012). Such data are also needed to further test for general consequences of asexuality beyond lineage-specific effects (Jaron *et al*. 2020). For many animal groups in which asexuality is frequent, genomic data are limited to a few representatives only (Tvedte *et al*. 2019) or are totally absent like in the Ostracoda.

Of all extant non-marine ostracods, the Cyprididae (cyprids) are most speciose, comprising 42% of all known species (Meisch *et al*. 2019). They would thus be an obvious choice for genomic studies, also because in this ostracod family, mixed reproduction with sexual and asexual females and geographic parthenogenesis is very common (Horne *et al*. 1998). Asexual cyprids, however, are often polyploid (Symonova *et al*. 2018; Adolfsson *et al*. 2010), probably because of hybridization between males and asexual females through accidental mating (Schmit *et al*. 2013). Consequently, genome sizes are relatively large (Jeffery *et al*. 2017; Gregory 2020) up to 3.13 pg which equals more than 3 Gb. These features are likely to seriously complicate genomic assemblies and annotations in the absence of any genomic resources for ostracods, which is why we did not choose any asexual cypridid ostracods for this genome project. Instead, we have selected three other species of nonmarine ostracods, one putative ancient asexual ostracod and two species with obligate sexual reproduction.

The ostracod family Darwinulidae is one of the two last remaining animal groups which are most likely true ancient asexuals (Heethoff *et al*. 2009; Schön *et al*. 2009b; Schwander 2016) and comprises about 35 morphospecies (Meisch *et al*. 2019). All darwinulids are brooders with valve dimorphisms between males and females that are detectable in the fossil record. Martens *et al*. (2003) showed that males have been absent in this family for at least 200 myr. One study reported a few males in a single darwinulid species (Smith *et al*. 2006) but proof of the functionality of these males for successful mating and meaningful genetic exchange could not been provided. Such (potential) atavistic males have also been reported in other putative ancient asexuals (Heethoff *et al*. 2009). The type species of the Darwinulidae, *Darwinula stevensoni*, has been asexual since c 20 myr (Straub 1952), occurs on all continents except Antarctica (Schön *et al*. 2012) and in a wide range of habitats (Schön *et al*. 2009b). *Darwinula stevensoni* is the best investigated darwinulid ostracod so far and has been the subject of ecological (Van Doninck *et al*. 2002, 2003a & b; Van den Broecke *et al*. 2013) and molecular research using DNA sequence data from single genes (Schön *et al*. 1998; Schön *et al*. 2003; Martens *et al*. 2005; Schön *et al*. 2012). These studies revealed that *D. stevensoni* is most likely apomictic or functionally mitotic (following the definition of apomixis in animals as in Schön *et al*. 2009a). The species also has low mutation rates. as there appears to be no (Schön *et al*. 1998) or low (Schön and Martens 2003; Schön *et al*. 2009b) allelic divergence within individuals, and genetic differences between populations from different continents can be attributed to ancient vicariant processes (Schön *et al*. 2012). It has also been suggested that gene conversion is common in this species (Schön and Martens 1998, 2003). These results, however, were based on a limited number of genes and require further confirmation with genome-wide data. *Darwinula stevensoni* has a life cycle of 1 year in Belgium (Van Doninck *et al*. 2003b) and up to 4 years in more northern regions (McGregor 1969 in Northern America; Ranta 1979 in Finland), which is exceptionally long for a non-marine ostracod. It can survive a wide range of temperatures, salinities (Van Doninck *et al*. 2002) and oxygen concentrations (Rossi *et al*. 2002). The total genome size of *D. stevensoni* has been estimated as 0.86-0.93 pg with flow cytometry (Paczesniak, unpublished), approximating 900 Mb. There is no information on the ploidy level of *D. stevensoni*, except for the study by Tétart (1979) showing 22 dot-like chromosomes. Because of its putative ancient asexuality, no close sexual relatives of *D. stevensoni* are available for comparative, genomic analyses. We have chosen two fully sexual non-marine ostracod species from the Cytherideidae and the Notodromadidae with high population densities in Belgium as comparisons to the putative ancient asexual: *Cyprideis torosa* and *Notodromas monacha* respectively. *Cyprideis torosa* inhabits brackish waters and is the only extant species of this genus in Europe (Meisch 2000). It has been the subject of various biological and especially palaeontological and geochemical studies (see for example: Heip, 1976a, b; De Deckker *et al*. 1999; Keyser 2005). Frogley and Whittacker (2016) suggested that *C. torosa* is at least of Pleistocene origin (c 2.5 myr) but might be older. There are only two molecular studies of this species based on single genes (Schön and Martens 2003; Schön *et al*. 2017). No information on the genome size or the karyotype of *C. torosa* is currently available.

The second sexual ostracod species analysed here, *Notodromas monacha*, occurs throughout the Northern hemisphere and is a non-marine ostracod with a most peculiar behaviour: it is partially hyponeustonic, hanging upside down attached to the water surface (Meisch 2000). The fossil record of *N. monacha* goes back to the Miocene (max 23 myr - Janz 1997), and its genome size is at 0.87pg (Jeffery *et al*. 2017; Gregory 2020) very similar to that of *D. stevensoni*. This species has not yet been the subject of any molecular studies.

Our aim here is to provide the first reference genome data of non-marine ostracods from three different species with varying reproductive modes: the putative ancient asexual *D. stevensoni* and the two obligate sexuals, *C. torosa* and *N. monacha*. We also generate transcriptomes of these species to facilitate genome annotations.

## MATERIAL AND METHODS

### Sample collection for genome and transcriptome sequencing

All three non-marine ostracod species were sampled in Belgian lakes where previous research had shown that these species occurred (Schön and Martens 2003; Merckx *et al*. 2018). Living ostracods were sampled using a hand net with a mesh size of 150 μm. The hand net was swept in between the vegetation and forcefully right above the surface of the sediment for collecting *Darwinula stevensoni* and *Cyprideis torosa*. *Notodromas monacha* was sampled by moving the net on the water surface. Non-marine ostracods were kept in habitat water. Their taxonomic identity was confirmed, and they were sorted alive under a binocular microscope as described by Martens and Horne (2016). Individual ostracods were picked with a pipette and transferred into sterilized EPA water in which they were maintained until DNA and RNA was extracted. More details on the origin of biological samples are provided in Table S2.

For generating reference genomes, DNA was extracted from a single female of each species using the QIAamp DNA Micro kit according to manufacturer’s instructions. The extracted DNA from single females was amplified in two independent reactions using the SYNGIS TruePrime WGA kit and then pooled, to generate sufficient DNA for preparing different libraries. To generate transcriptomes for annotation of reference genomes, RNA was extracted from 40 pooled individuals per species from the same collection batch. For this, individuals were frozen in liquid nitrogen and, after addition of Trizol (Life Technologies), mechanically crushed with beads (Sigmund Lindner). Next, chloroform and ethanol-treatment was applied to the homogenized tissue and the aqueous layer transferred to RNeasy MinElute Columns (Qiagen). Subsequent steps of RNA extraction were done following the RNeasy Mini Kit protocol, including DNase digestion. Finally, RNA was eluted into water and stored at −80°C. RNA quantity and quality were estimated with the NanoDrop (Thermo Scientific) and Bioanalyzer (Agilent).

### Genome assembly

We prepared five genomic DNA libraries for each reference genome (three 2×125bp paired-end libraries with average insert sizes of 250-300, 550 and 700bp, and two mate-pair libraries with average insert sizes of 3000 and 5000bp; see Table S3 for more details) with the Illumina TruSeq DNA Library Prep Kit. Reads were generated with the Illumina HiSeq 3000 system for a total coverage between 351X and 386X (Table S3).

Reads were filtered with Trimmomatic v0.36 (Bolger *et al*. 2014) and NxTrim v0.4.1 (O’Connell *et al*. 2015). We employed non-standard methods for *de novo* genome assemblies owing to uneven coverage produced by PCR-based whole-genome amplification (Chen *et al*. 2013; Oyola *et al*. 2014)._Filtered reads were normalized using BBMap v36.59 (Bushnell 2014) and then assembled into contigs with SPAdes v3.10.1 (Bankevich *et al*. 2012). Scaffolding was performed using SSPACE v3.0 (Boetzer *et al*. 2012). Scaffolds identified as contaminants were filtered out using Blobtools v1.0 (Laetsch and Blaxter 2017). The completeness of genomes assemblies was assessed with BUSCO v3.0.2 (Seppey *et al*. 2019) against the *arthropoda_odb9* lineage. More details of the assembly pipelines and the applied parameters can be found in supplementary materials (SM1).

### Protein coding gene annotation

Libraries were prepared using the Illumina TruSeq Stranded RNA, following the manufacturer’s instructions. RNA reads were generated with the Illumina HiSeq 2500 system (Table S4). Reads were filtered with Trimmomatic v0.36. All trimmed reads were mapped against the genomes with STAR v2.5.3a (Dobin *et al*. 2013) and further assembled with Trinity v2.5.1 (Haas *et al*. 2013) under the “genome guided” mode to produce transcriptome assemblies.

The obtained transcriptomes and protein evidence were used to train and predict protein coding genes using MAKER v2.31.8 (Holt and Yandell 2011). Predicted protein coding genes were functionally annotated with Blast2GO v5.5.1 (Conesa *et al*. 2005; Götz *et al*. 2008) against the NCBI *non-redundant arthropods* protein database (v 2018-10). More details of the annotation pipelines and the applied parameters can be found in supplementary materials (SM2).

### GenomeScope analyses

The whole genome amplification approach, which we used in the present study because of the small body size of individual ostracods, generated unequal read coverage of ostracod genomes and prevented us from directly estimating genome sizes and levels of heterozygosity from the assemblies. To overcome this problem, we re-sequenced two individual ostracods each of *D. stevensoni* and *N. monacha* without whole genome amplification, preparing libraries with the NEBNext^®^ Ultra™ II DNA Library Prep Kit for Illumina. Reads were filtered with Trimmomatic v0.36 and analyzed using GenomeScope v2.0 (Ranallo-Benavidez *et al*. 2020) to correctly estimate genome size and heterozygosity. More details on the analyses are provided in the supplementary material (SM3).

## RESULTS AND DISCUSSION

### First ostracod reference genomes and their features

We successfully produced the first *de novo* reference genomes of non-marine ostracods, namely of the three species *Darwinula stevensoni, Cyprideis torosa* and *Notodromas monacha* with different reproductive modes (see SM1 and Tables S3–S4 for more details on the assemblies). Given the small size of individual non-marine ostracods and the limited amount of soft tissue suitable for DNA and RNA extractions (Schön and Martens 2016), this was not a trivial task. We used a whole genome amplification approach (WGA), because the TruSeq DNA Nano library prep kit for Illumina sequencing or low input protocols for PacBio (Duncan *et al*. 2019) were not yet available when these assemblies were generated. We would not recommend WGA for future studies because this PCR-based method generated uneven coverage, and consequently, problems for applying routine genome assembly methods and estimates of genome size and heterozygosity. Despite these limitations, our approach produced genome assemblies that are useful for future research as will be outlined below.

When assessing the quality of the obtained ostracod *de novo* genome assemblies, the assembly of the putative ancient asexual, *D. stevensoni*, had the best contiguity, with the largest N50 although the total number of scaffolds was similar to *N. monacha* (Table 1, Table S6). The genome of the putative ancient asexual is furthermore the most complete as shown by its total BUSCO score of 96% and of 94% for complete single copy genes (Table 1). The quality of the genome from the obligate sexual ostracod *Cyprideis torosa* is the lowest of the three ostracod species as it has the highest number of scaffolds, and the lowest N50; it is also less complete with a total BUSCO score of 92% (Table S7) and of 87% for complete single copy genes (Table 1). All three species have similar numbers of predicted genes and transcripts (Table S7).

**Table 1:** Quality features of published crustacean genomic assemblies of the last four years and of the current study. Assembly size is provided in million base pairs, scaffold N50 in kilo base pairs and BUSCO scores in %. Letters behind BUSCO scores indicate the % of complete single copy genes (C) or % of single and fragmented single copy genes (C+F), respectively. Where BUSCO scores lack brackets, no further information on completeness of single copy genes was provided. $ anchoring of scaffolds in existing genome assembly; L = linkage map available; * long read technology; # BAC library available. n. i. = no information available.

| Class | Order | Species | Size | # scaffolds | N50 | BUSCO | Reference |
| --- | --- | --- | --- | --- | --- | --- | --- |
| Branchiopoda | Diplostraca | *Daphnia pulex*<sup>SL</sup> | 156 | 1,822 | 1,661 | 96 | Ye *et al.* 2017 |
| Branchiopoda | Diplostraca | *D. magna*<sup>SL</sup> | 130 | 4,193 | 10,124 | 96.7 (C) | Lee *et al.* 2019 |
| Branchiopoda | Notostraca | *Lepidurus arcticus* | 73 | 7,167 | 116 | 98.4 (C) | Savojardo *et al.* 2018 |
| Branchiopoda | Notostraca | *L. apus lubbocki* | 90 | 20,738 | 402 | 97.8 (C) | Savojardo *et al.* 2018 |
| Branchiopoda | Spinicaudata | *Eulimnadia texana* * | 120 | 112 | 18,000 | n. i. | Baldwin-Brown *et al.* 2017 |
| Copepoda | Cyclopoida | *Apocyclops royi* | 258 | 97,072 | n. i. | 50 (C) | Jørgensen *et al.* 2019 |
| Copepoda | Cyclopoida | *Oithona nana* | 85 | 4,626 | 401 | n. i. | Madoui *et al.* 2017 |
| Copepoda | Harpaticoida | *Tigriopus californicus* * | 190 | 459 | 298 | 94.5 (C) | Barreto *et al.* 2018 |
| Copepoda | Harpaticoida | *T. japonicus*<sup>\$</sup> | 197 | 339 | 10,650 | 96 (C) | Jeong *et al.* 2020 |
| Copepoda | Harpaticoida | *T. kingsejongensis* | 295 | 270,823 | 159 | 61.1 (C) | Kang *et al.* 2017 |
| **Ostracoda** | **Podocopida** | ***Cyprideis torosa*** | **335** | **132,611** | **19** | **86.6 (C)**<br>**91.9 (C+F)** | **Current study** |
| **Ostracoda** | **Podocopida** | ***Darwinula stevensoni*** | **382** | **62,118** | **56** | **93.7 (C)**<br>**95.8 (C+F)** | **Current study** |
| **Ostracoda** | **Podocopida** | ***Notodromas monacha*** | **377** | **62,251** | **42** | **92.7 (C)**<br>**94.4 (C +F)** | **Current study** |
| Malacostraca | Amphipoda | *Parhyale hawaiiensis*<sup>#L</sup> | 4,024 | 100,000 | 69 | n. i. | Kao *et al.* 2016 |
| Malacostraca | Isopoda | *Armadillidium vulgare* * | 1,725 | 43,451 | 51 | 87.9 (C) | Chebbi *et al.* 2019 |
| Malacostraca | Decapoda | *Cherax quadricarinatus* * | 3,237 | 508,682 | 33 | 81.3 (C) | Tan *et al.* 2020 |
| Malacostraca | Decapoda | *Eriocheir japonica sinensis* * | 1,270 | 1,368 | 3,185 | 92.7 (C) | Tang *et al.* 2020 |
| Malacostraca | Decapoda | *Palaeomon carinicauda*<sup>L</sup> | 9,185 | 28,089,718 | 586 | n. i. | Li *et al.* 2019 |
| Malacostraca | Decapoda | *Penaeus monodon* * | 1,600 | 1,211,364 | 2 | 96.8 (C+F) | Van Quyen *et al.* 2020 |
| Malacostraca | Decapoda | *Litopenaeus vannamei*<sup>*L</sup> | 1,664 | 4,682 | 606 | 95 | Zhang *et al.* 2019 |
| Malacostraca | Decapoda | *Marsupenaeus japonicus* | 924 | 37,192,281 | 1 | 97 | Yuan *et al.* 2018 |
| Malacostraca | Decapoda | *Procambarus virginalis* | 3,300 | 3,752,011 | 39 | n. i. | Gutekunst *et al.* 2018 |

Ostracod genome sizes estimated with flow cytometry are somewhat larger than the estimates that we obtained here from GenomeScope analyses of re-sequenced individual ostracods. The haploid genome size of *D. stevensoni* was estimated at 420-455 Mb with flow cytometry (Paczesniak, unpublished) while we estimated 362 Mb from sequence reads (Figure S1 A-B). Similarly, the size of the haploid genome of *N. monacha* is estimated at 425 Mb with flow cytometry (Jeffery *et al*. 2017; Gregory 2020), which is larger than the 385 Mb (Figure S1 CD) that we obtained from sequence reads. It thus seems that some parts of each genome are missing from our sequencing reads. Transposons and repeat-rich genomic regions can contribute to gaps in genomic assemblies (Peona *et al*. 2020). Some of these missing regions could also be GC-rich, a feature which is known to cause a sequencing-bias with Illumina technology (see for example Chen *et al*. 2013, Botero-Castro *et al*. 2017). Acquiring more complete genome assemblies will require the additional application of long read technologies to ostracods.

Genome-wide estimates of heterozygosity are especially interesting for asexual taxa because the absence of recombination is expected to cause accumulation of mutations, resulting in increasing allelic divergences within individuals (Birky 1996). Jaron *et al*. (2020) identified three factors driving intragenomic heterozygosity in asexuals: how the transition to parthenogenesis occurred, which cytological mechanism underlies parthenogenesis and how long asexual reproduction has been ongoing. Based on sequencing reads from individual ostracods, we estimate heterozygosity of the putative ancient asexual ostracod *D. stevensoni* to be 0.92-0.99% (Figure S1 A-B) and 1.32-1.43% for the sexual *N. monacha* (Figure S1 CD). The genome-wide heterozygosity of *D. stevensoni* matches to some extent an earlier study on intra-individual divergence in three nuclear genes of *D. stevensoni* (Schön and Martens 2003). The finding of almost 1% heterozygosity in *D. stevensoni* is remarkable, given that all previous genome-wide estimates for asexual arthropods that did not evolve via hybridization revealed extremely low levels of heterozygosity (Jaron *et al*. 2020). Yet heterozygosity is clearly less than the estimates for parthenogenetic species with known hybrid origin (1.73-8.5%) or polyploidy (1.84%-33.21%) (Jaron *et al*. (2020), supporting the view that *D. stevensoni* is neither a hybrid nor a polyploid. Asexual reproduction in ostracods is thought to be apomictic (Chaplin *et al*. 1994), implying that observed heterozygosity levels are largely depend on the relative impact of heterozygosity losses from gene conversion and heterozygosity gains from new mutations. Given the apparent absence of sex and recombination for millions of years (Straub 1952), it is perhaps surprising that heterozygosity in this putative ancient asexual ostracod is not larger. This may suggest that genome-wide rates of gene conversion and mutation are comparable in this species.

### Genome contiguity of ostracod assemblies as compared to other crustaceans

We here compare the qualities of our ostracod genome assemblies to those of 19 other crustacean species (Table 1) published in the last four years. We only include studies with complete assemblies and sufficient information to assess assembly qualities. We assessed the contiguity of the three *de novo* ostracod genome assemblies by the number of scaffolds and their N50. On the one hand, both features are comparable to those of the copepod *Apocylops royi* (Jørgensen *et al*. 2019) and the amphipod *Parhyale hawaiensis* (Kao *et al*. 2016) (Table 1) and better than for crustaceans with larger genomes such as the decapods *Cherax quadricarinatus* (Tan *et al*. 2019), *Palaeomon carinicauda* (Li *et al*. 2019), *Penaeus mondon* (Van Quyen *et al*. 2020), *Marsupenaeus japonicus* (Yuan *et al*. 2018) and *Procamburus virginalis* (Gutekunst *et al*. 2018; Table 1). On the other hand, genome assemblies of several other crustaceans have smaller scaffold numbers and higher N50 and thus better contiguities than the assemblies obtained here for non-marine ostracods. For the two notostracan *Lepidurus* species (Savojardo *et al*. 2018), this can probably be explained by their smaller genome sizes. For other crustaceans, genome assemblies or linkage maps have been available beforehand which have considerably improved assembly qualities (Table 1) as in the examples of the cladocerans *Daphnia pulex* (Ye *et al*. 2017), *D. magna* (Lee *et al*. 2019) and the copepod *Tigriopus japonicus* (Jeong *et al*. 2020). No such genomic resources are currently available for ostracods. Finally, other studies of crustacean genomes with better assembly contiguities (the branchiopod *Eulimnadia texana* - Baldwin-Brown *et al*. 2018, and the decapod *Erichoir japonica sinensis* - Tang *et al*. 2020), the copepod *Tigriopus californicus* - Jeong *et al*. 2020 and the isopod *Armadillium vulgare* - Chebbi *et al*. 2019) have used a combination of Illumina and long read technologies (Table 1). Long-read technologies such as PacBio used to require a relatively large amount of high-molecular weight DNA (Solares *et al*. 2018), which could not be obtained for ostracods with their very low yields of high molecular weight DNA from individual specimens and their small body sizes as compared to many other crustaceans (Schön and Martens 2016). We hope that low input protocols for PacBio (Duncan *et al*. 2019) and other long read technologies can be successfully applied to ostracods in the future, in which case the genome assemblies obtained here could form the basis for subsequent hybrid assemblies. Optimizing Oxford Nanopore Technology for non-marine ostracods has already commenced (Schön et al. in prep.).

### Genome annotations of ostracods and other crustaceans

Because our *de novo* ostracod genome assemblies are relatively complete (see BUSCO scores in Table 1), we will here also briefly compare some features of predicted protein coding genes with those of other crustaceans (Table S8). We have predicted 13,771 to 17,776 protein coding genes in the three non-marine ostracod genomes, (Tables S7 and S8) with the highest number for the sexual *C. torosa* and an intermediate estimate for the putative ancient asexual *D. stevensoni*. The number of annotated protein coding genes in non-marine ostracods is similar to estimates for various branchiopods and the copepods *Oithona nana, Tigriopus californicus* and *T. kingsejongensis* but lower than in most malacostracans (Table S8). Not all genome studies of crustaceans cited here contain information on other features of coding genes, such as the average size of genes, introns and exons (Table S7). Comparisons of these features are therefore limited and will not be further discussed here but we provide available data of these features for ostracods and other crustacean genomes for reference.

Gene annotation in general but especially in the crustaceans is challenging; this is for example illustrated by the much lower numbers of protein coding genes (18,440) which are predicted in the novel reference genome of the cladoceran *Daphnia pulex* by Ye *et al*. (2017) as compared to the first assembly of *D. pulex* with more than 30,000 predicted genes (Colbourne *et al*. 2011). Even more difficult is assigning gene functions to annotated crustacean genomes (Rotlantt *et al*. 2018). The novel data on predicted genes and transcripts from non-marine ostracods in the current study will significantly contribute to future genome annotations in crustaceans and other arthropods. The genes and transcripts predicted here can also provide the baseline for future gene expression studies of non-marine and marine ostracods.

## CONCLUSIONS

We have successfully obtained *de novo* genome assemblies for three species of non-marine ostracods with different reproductive modes. These represent the first quality reference genomes for ostracods. Given the paucity of genome assemblies from crustaceans as compared to insects or other arthropods, these assemblies are important tools to further develop ostracods as models for evolutionary and ecological research, also including marine species. Even if the *de novo* genome assemblies are somewhat fragmented and not yet at the chromosome-level, they have a high level of completeness and will thus facilitate future studies of ostracods. The reference genomes of this paper can also provide the first step towards a genomic assessment of the putative ancient asexual status of non-marine darwinulid ostracod species.

## DATA AVAILABILITY

Raw sequence reads have been deposited in NCBI’s sequence read archive under the following bioprojects: PRJNA515625 (reference genomes, Table S3) and PRJNA631617 (RNA-seq for annotations and resequenced individuals, Table S4 and Table S5).

Genome assemblies and annotations have been deposited in the European Nucleotide Archive (ENA) under the accession number PRJEB38362 (Table S6 and Table S7). Codes for the analyses are available at: https://github.com/AsexGenomeEvol/Ostracoda_genomes.

## ACKNOWLEDGEMENTS, FUNDING, CONFLICT OF INTEREST

This research was funded by Belgian Federal Science Policy (BR/314/PI/LATTECO) and a grant from the Swiss National Science Foundation (CRSII3_160723). Marie Cours, Tijs Van Den Bergen and Jeroen Venderickx are acknowledged for technical support in sampling and sorting ostracod samples. We also thank Kristiaan Hoedemakers and Jeroen Venderickx for their assistance in finalizing the figure.

## Supplementary material

### SM1: Details of genome assembly pipelines and parameters

The paired-end reads were filtered using Trimmomatic v0.36 (Bolger *et al*. 2014). Adapters were removed using adapters provided by Trimmomatic. Leading and trailing bases below quality 3 were removed. Reads were scanned using a 4-base sliding window and trimmed when the average quality dropped below 20. Reads were discarded if the size dropped below 60 bp. (parameters: PE ILLUMINACLIP:TruSeq-PE.fa:2:30:10:4 LEADING:3 TRAILING:3 SLIDINGWINDOW:4:15 AVGQUAL:20 MINLEN:60).

The mate-pair reads were filtered using NxTrim v0.4.1 (O’Connell *et al*. 2015) (parameters: --preserve-mp --separate). As a result, mate-pair reads flagged as MP and UNKNOWN were concatenated following the authors’ suggestion. Finally, the concatenated mate-pair reads were trimmed using Trimmomatic with the same parameters described previously.

We employed non-standard methods for genome assemblies due to uneven coverage produced by PCR-based whole-genome amplification (Chen *et al*. 2013; Oyola *et al*. 2014). Only the overlapped libraries were used for contig assemblies. Filtered paired-end reads were merged into single reads by overlap detection using BBMap v36.59 (Bushnell 2014) and the module BBMerge (parameters: minoverlap=15 mismatches=0 ecct strict). The merged reads were then concatenated with the rest of the single reads and normalization was performed using the module BBNorm with a target average depth of 65x. Normalized data were *de novo* assembled using SPAdes v3.10.1 (Bankevich *et al*. 2012) (parameters: --careful -k 21,33,55,77,99,111,127). Contigs were then scaffolded with SSPACE v3.0 (Boetzer *et al*. 2012) using the remaining libraries and gap-filled with GapCloser v1.12-r6, a module of the SOAP package (Luo *et al*. 2012) with default parameters. Then, the genome assemblies were decontaminated using BlobTools v1.0 (Laetsch & Blaxter, 2017) under the taxrule “bestsumorder”. Hit files were generated after a blastn v2.7.1+ against the NBCI nt database (v 2016-06), searching for hits with an evalue below 1e-25 (parameters: -max_target_seqs 10 -max_hsps 1 -evalue 1e-25). Scaffolds without hits to metazoans were filtered out from the assemblies.

The completeness of genomes assemblies was assessed with BUSCO v3.0.2 (Seppey *et al*. 2019) against the *arthropoda_odb9* lineage and the --long option.

### SM2: Annotation of protein coding genes

Raw paired-end RNA-sequence reads were trimmed according to quality with Trimmomatic v0.36 (Bolger *et al*. 2014) using the maximum information approach with tolerant parameters.

Any reads with a sequence length of < 80 bp after trimming were discarded (parameters: adapters.fa:2:30:12:1:true LEADING:3 TRAILING:3 MAXINFO:40:0.4 MINLEN:80). All trimmed RNA-sequence reads were then mapped against the genomes using STAR v2.5.3a (Dobin *et al*. 2013) under the “2-pass mapping” mode and default parameters. Then, the STAR outputs were used to produce transcriptome assemblies using Trinity v2.5.1 (Haas *et al*. 2013) “genome guided” mode (parameters: -- genome_guided_max_intron 100000 --SS_lib_type RF --jaccard_clip). Finally, the transcriptome assemblies were filtered following Trinity developers recommendations (https://github.com/trinityrnaseq/trinityrnaseq/wiki/Trinity-FAQ): Briefly, filtered RNA-seq reads were mapped back against the transcriptomes using Kallisto v0.43.1 (Bray *et al*. 2016) with options --bias and --rf-stranded then transcripts with at least 1 TPM in any samples were retained.

Protein coding genes were predicted using MAKER v2.31.8 (Holt & Yandell 2011) in a 2-iterative way described in Campbell *et al*. (2014) with minor modifications following author recommendations. Only scaffolds above 500bp were annotated. Prior to gene prediction, MAKER used RepeatMasker v4.0.7 (Tarailo-Graovac & Chen 2009) for masking repetitive regions. For the first iteration, genes were predicted using Augustus v3.2.3 (Stanke *et al*. 2006) trained with the BUSCO v3.0.2 (Seppey *et al*. 2019) results (SM1). A combination of UniProtKB/Swiss-Prot (release 2018_01) and the BUSCO *arthropoda_odb9* proteome were used as protein evidence. The Trinity assembled mRNA-seq reads (described above) were used as transcript evidence. The resulting gene models were then used to retrain Augustus as well as SNAP v2013.11.29 (Korf 2004) and a second iteration was performed. Subsequently, predicted protein coding genes were functionally annotated using Blast2GO v5.5.1 (Conesa *et al*. 2005; Götz *et al*. 2008) with default parameters against the NCBI *non-redundant arthropods* protein database (v 2018-10).

### SM3: Estimation of genome size and heterozygosity by genome profiling analysis

Raw paired-end resequencing reads were trimmed using the same strategy as described in SM2. K-mer frequencies were computed using KMC v3.1.1 (Kokot *et al*. 2017). Genome sizes and heterozygosities were estimated with GenomeScope v2.0 (Ranallo-Benavidez *et al*. 2020) using parameters recommended by the authors.

**Table S1:** Overview of RNA sequencing reads from ostracods in GenBank. No full assemblies or annotations are available from these studies. # of genes = total number of orthologous genes used for phylogenetic or gene expression studies, respectively.

| Order | Species | GenBank accession number | Sequencing technique | # of genes | Study purpose | Reference |
| --- | --- | --- | --- | --- | --- | --- |
| Mydocopida | *Vargula tsujii* | SRX532406 | 454 | 261 | Phylogeny of pancrustaceans | Oakley *et al.* 2012 |
| Mydocopida | *Euphilomedes morini* | SRX1097309 | 454 | 261 | Phylogeny of pancrustaceans | Oakley *et al.* 2012 |
| Mydocopida | *Alternochelata lizardensis* | PRJNA436031 | Illumina |  | 1KITE project | unpublished |
| Mydocopida | *Vargula hilgendorfii* | SRX884494 | Illumina |  | 1KITE project | unpublished |
| Mydocopida | *Vargula hilgendorfii* | DRX059606 | Illumina |  |  | unpublished |
| Mydocopida | *Eusarsiella sp.* | SRX2085852 | Illumina | 244 | Phylogeny of pancrustaceans | Lozano-Fernandez *et al.* 2019 |
| Mydocopida | *Cylindroleberidinae* | SRX2085850 | Illumina | 244 | Phylogeny of pancrustaceans | Lozano-Fernandez *et al.* 2019 |
| Mydocopida | *Rutiderma sp.* | SRX2458829 | Illumina | 277 | Phylogeny of pancrustaceans | Schwenter *et al.* 2018 |
| Halocyprida | *Conchoecia obtusata* | SRX2458823 | Illumina | 277 | Phylogeny of pancrustaceans | Schwenter *et al.* 2018 |
| Podocopida | Undetermined; family Cypridinae | JL207200 | 454 | 822 | Phylogeny of pancrustaceans | Von Reumont *et al.* 2012 |
| Podocopida | *Heterocypris incongruens* | ICLE01000001 | Illumina | 4 | Expression of antioxidant genes | Hiki *et al.* 2019 |
| Podocopida | *Pontocypris mytiloides* | SRX4048873 | Illumina | 277 |  | unpublished |
| Podocopida | *Paranesidea sp.* | SRX2085851 | Illumina | 277 | Phylogeny of pancrustaceans | Schwenter *et al.* 2018 |
| Podocopida | *Pterygocythereis sp.* | SRX2458816 | Illumina | 277 | Phylogeny of pancrustaceans | Schwenter *et al.* 2018 |

**Table S2:** Origin of biological material. All species were collected in Belgium in 2018. The species printed in blue is a putative ancient asexual, species printed in red are sexually reproducing.

| Species | Population name | Habitat |
| --- | --- | --- |
| *Cyprideis torosa* | Dievengat | Natural brackish lake |
| *Darwinula stevensoni* | Zaventem | Artificial lake |
| *Notodromas monacha* | Overijse | Artificial lake |

**Table S3:** Statistics of ostracod genome sequence data. bp = basepairs. G = giga. The coverage is estimated from final assembly sizes. The species printed in blue is a putative ancient asexual, species printed in red are sexually reproducing. Coverage is provided in %.

| Species | Data type | Insert size (bp) | Yield (Gbp) | Coverage |
| --- | --- | --- | --- | --- |
| *Cyprideis torosa* | Paired-end 2x150 bp | 250-300 | 45.8 | 136.9 |
|  |  | 350 | 41.3 | 123.3 |
|  |  | 550 | 16.2 | 48.5 |
|  | Mate-pair 2x150 bp | 3000 | 13.6 | 40.4 |
|  |  | 5000 | 12.2 | 36.5 |
| *Darwinula stevensoni* | Paired-end 2x150 bp | 250-300 | 51.8 | 135.5 |
|  |  | 350 | 39.1 | 102.2 |
|  |  | 550 | 13.0 | 34.0 |
|  | Mate-pair 2x150 bp | 3000 | 18.7 | 48.9 |
|  |  | 5000 | 12.2 | 32.0 |
| *Notodromas monacha* | Paired-end 2x150 bp | 250-300 | 36.9 | 98.0 |
|  |  | 350 | 51.3 | 136.2 |
|  |  | 550 | 15.0 | 39.9 |
|  | Mate-pair 2x150 bp | 3000 | 15.5 | 41.1 |
|  |  | 5000 | 13.4 | 35.6 |

**Table S4:** Statistics of ostracod transcriptome sequence data. Data type: paired-end 2x 101bp. bp = basepairs. G = giga. The species printed in blue is a putative ancient asexual, species printed in red reproduce sexually.

| Species | Description | Yield (Gbp) |
| --- | --- | --- |
| *Cyprideis torosa* | Pool 1 (40 males and females) | 7.2 |
|  | Pool 2 (40 males and females) | 8.2 |
|  | Pool 3 (40 males and females) | 10.1 |
|  | Pool 4 (40 males and females) | 9.8 |
| *Darwinula stevensoni* | Pool 1 (40 females) | 6.7 |
|  | Pool 2 (40 females) | 7.1 |
|  | Pool 3 (40 females) | 7.4 |
| *Notodromas monacha* | Pool 1 (40 females) | 3.8 |
|  | Pool 2 (40 males) | 20.2 |

**Table S5:** Statistics of ostracod resequencing data. Data type: paired-end 2x 101bp. bp = basepairs. G = giga. The coverage is estimated from final assembly sizes. The species in blue is a putative ancient asexual, species printed in red reproduce sexually.

| Species | Description | Yield (Gbp) | Coverage |
| --- | --- | --- | --- |
| *Darwinula stevensoni* | Female DS_02 | 21.7 | 56.7 |
|  | Female DS_05 | 23.9 | 62.5 |
| *Notodromas monacha* | Female Nm_F03 | 20.9 | 55.6 |
|  | Female Nm_F07 | 24.0 | 63.8 |

**Table S6:** Statistics of genome assemblies of ostracod species. **Σ** represents the sum of all scaffolds in million basepairs (Mbp). The average N50 was calculated per scaffold in kilo basepairs (Kbp). The BUSCO score is the proportion of conserved single copy ortholog genes among arthropods. N is the proportion of unknown nucleotides (gaps) in the assembly. The species in blue is a putative ancient asexual, species printed in red reproduce sexually.

| Species | $\Sigma$<br>[Mbp] | N50<br>[kbp] | BUSCO<br>[%] | Ns<br>[%] |
| --- | --- | --- | --- | --- |
| *Cyprideis torosa* | 334.9 | 19.0 | 91.8 | 1.3 |
| *Darwinula stevensoni* | 382.1 | 56.4 | 95.8 | 0.9 |
| *Notodromas monacha* | 376.7 | 42.3 | 94.4 | 1.6 |

**Table S7:** Statistics of protein coding gene annotations and other gene features in ostracod genomes. bp = basepairs. The genome coverage is the proportion of each genome covered by genes. Red species reproduce sexually, while the species indicated in blue is a putative ancient asexual.

| Species | Gene count | Transcript count | Average gene length [bp] | Genome coverage [%] |
| --- | --- | --- | --- | --- |
| *Cyprideis torosa* | 17776 | 18069 | 4734 | 25.1 |
| *Darwinula stevensoni* | 15453 | 15922 | 7558 | 30.6 |
| *Notodromas monacha* | 13771 | 14294 | 7040 | 25.7 |

**Table S8:**
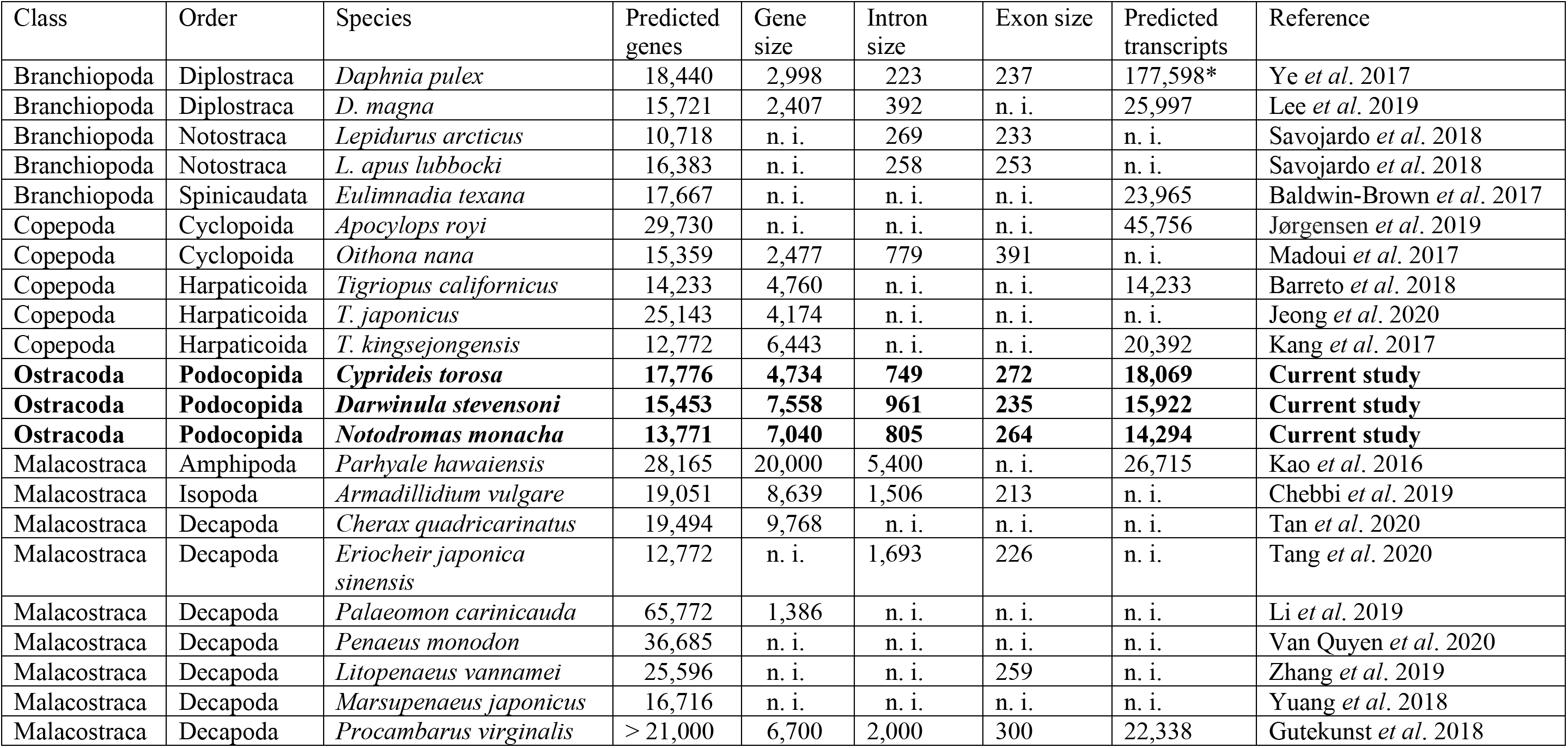
Annotations and gene features of crustacean genomes of the last four years and of the current study. Average gene, intron and exon sizes are provided in base pairs.* including all transcripts from different experiments and life stages. n. i. = no information.

**Figure S1.**
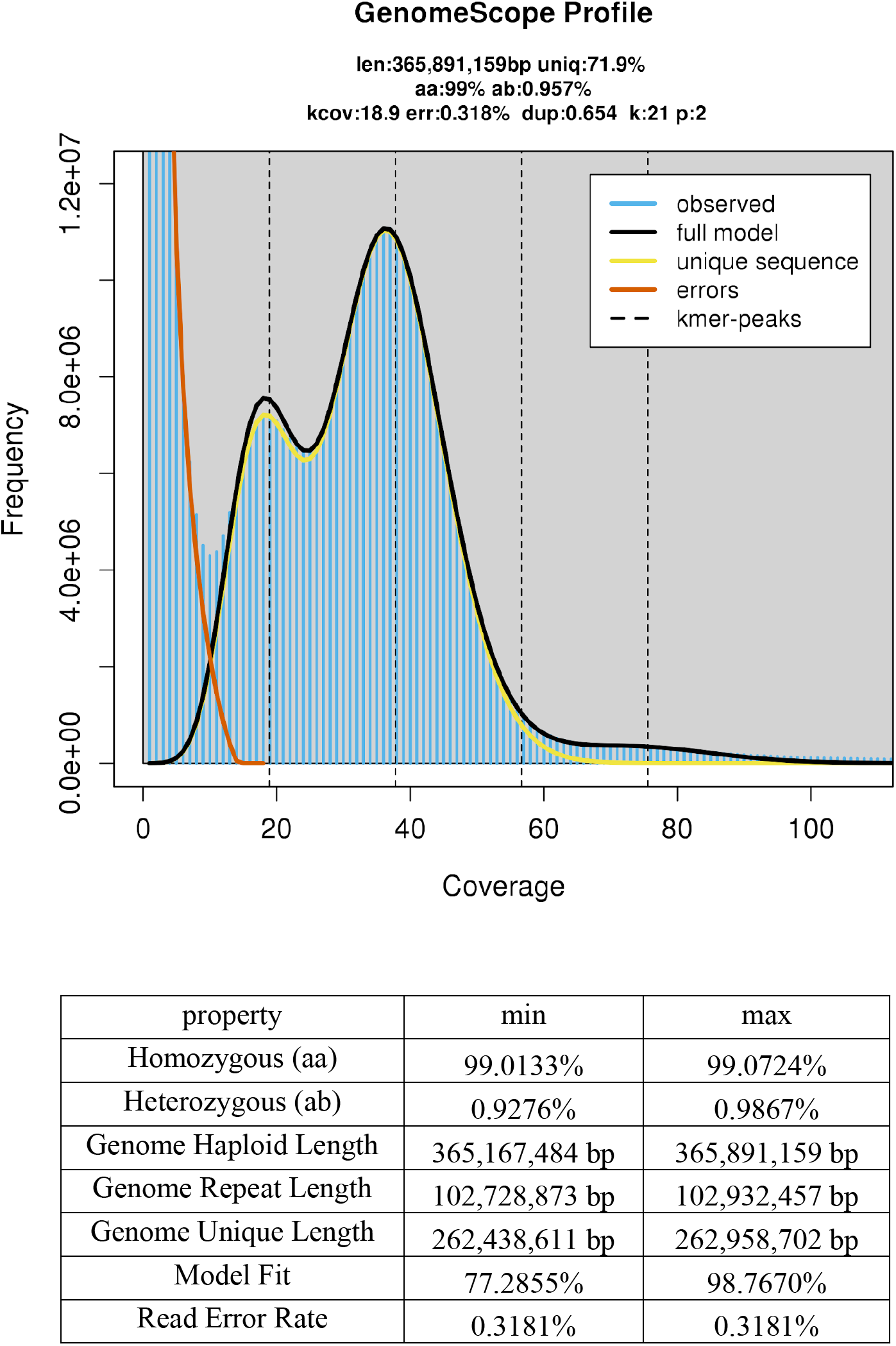
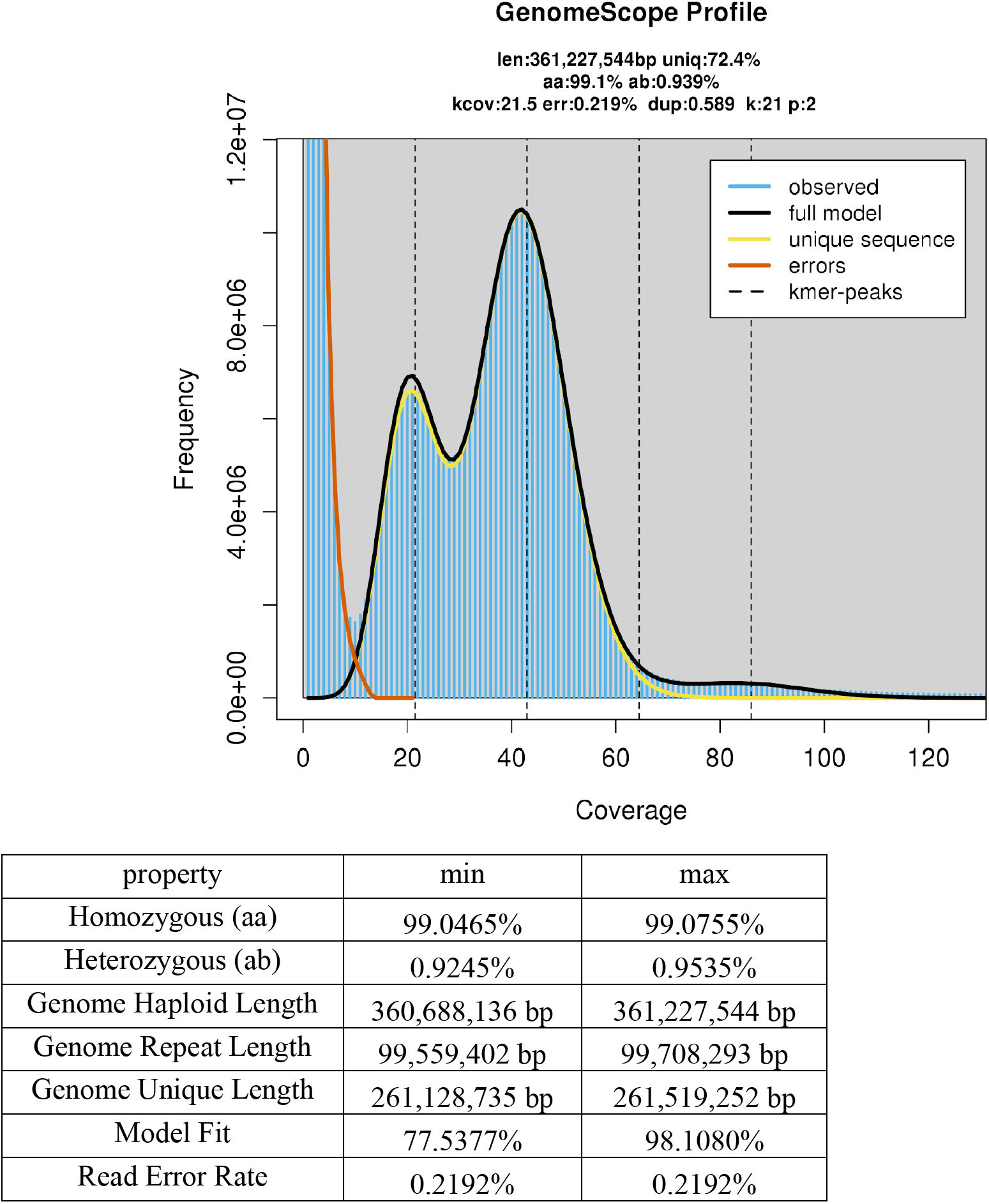
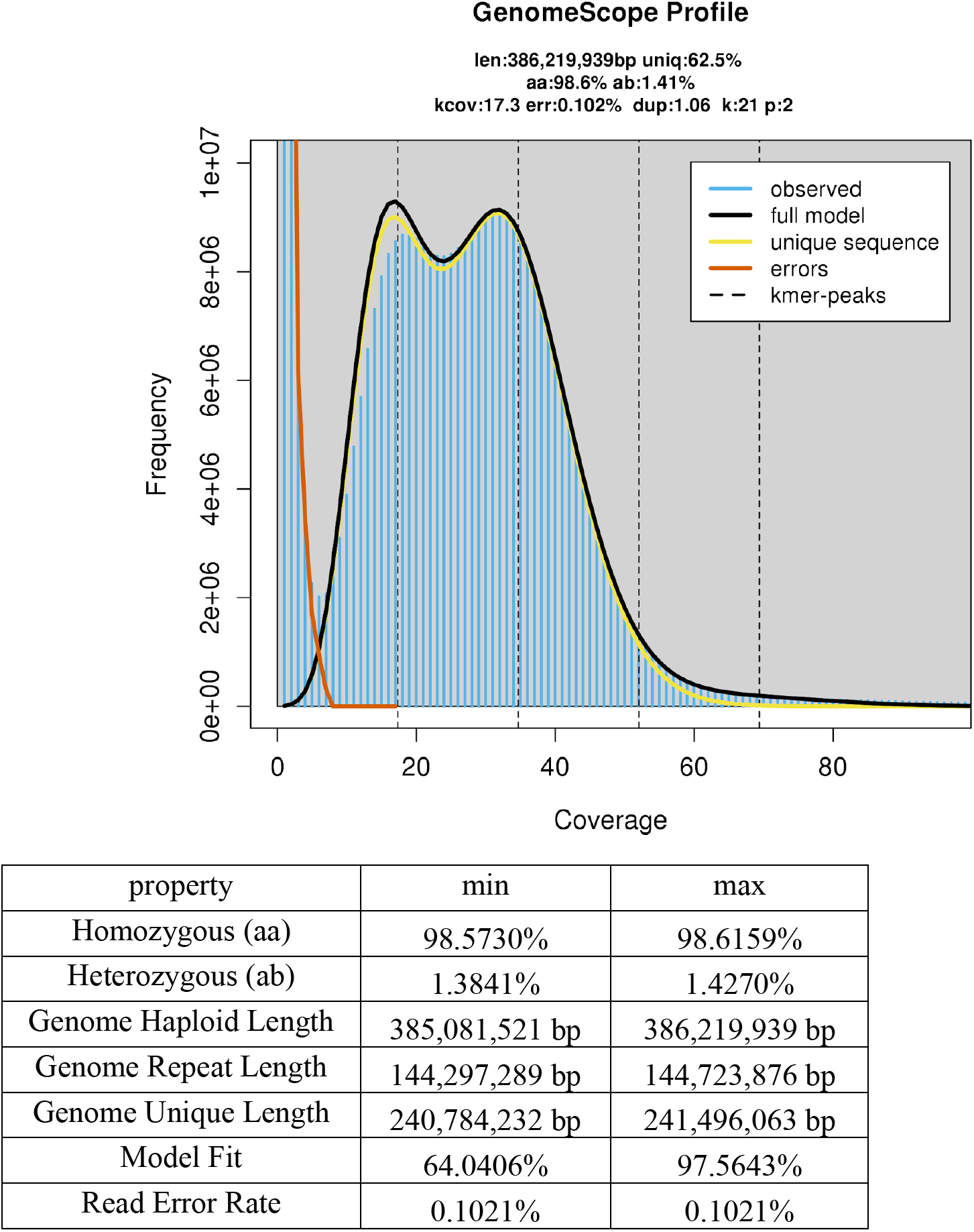
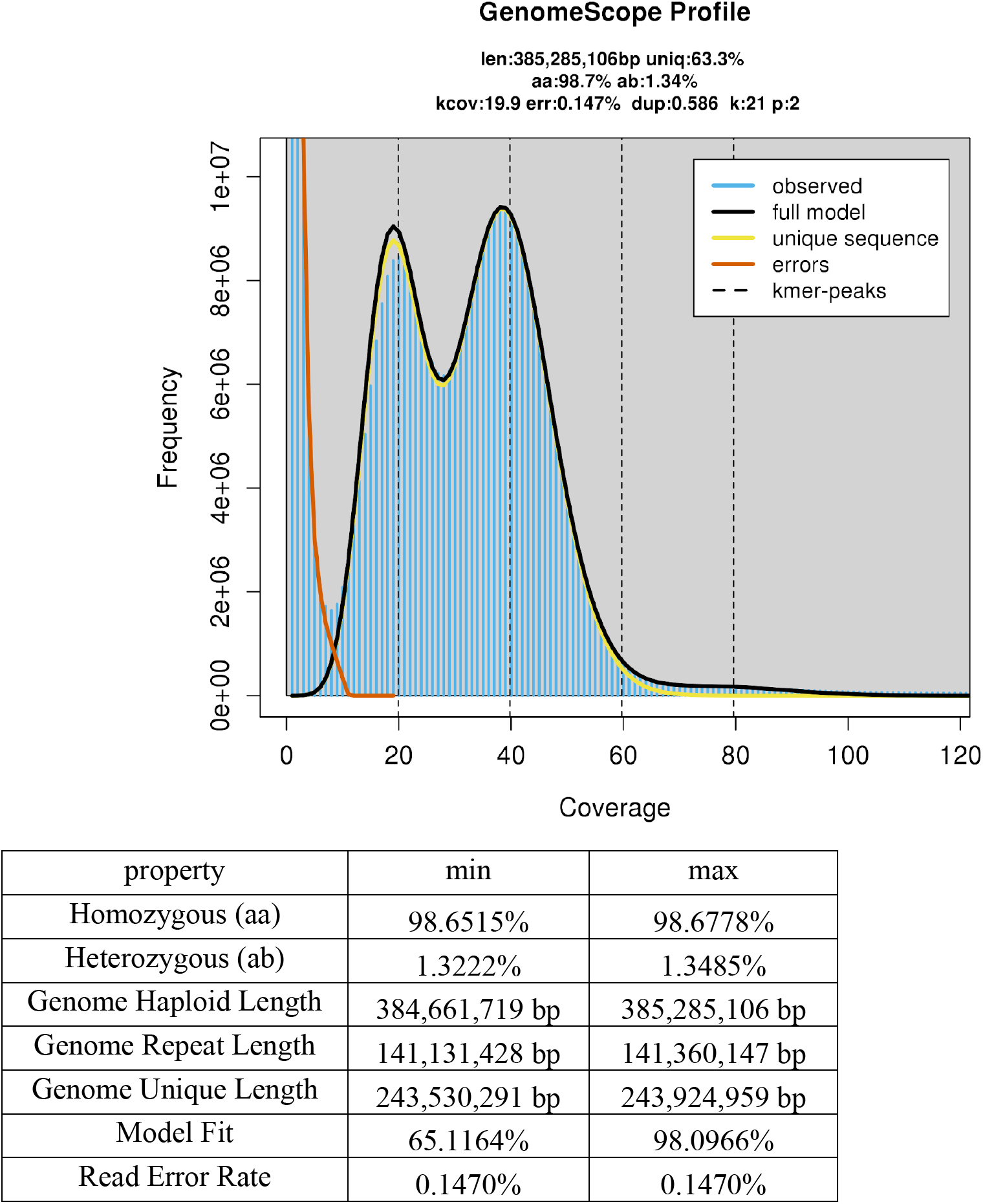
A-D: Results of GenomeScope analyses. Two individuals each of *Darwinula stevensoni* (Figure S1 A-B) and *Notodromas monacha* (Figure S1 CD) were resequenced and their reads analyzed to estimate genome sizes and heterozygosities. Figure S1 A – DS_02 Figure S1 B – DS_05 Figure S1 C – Nm_F03 Figure S1 D – Nm_F07

